# FGF10 triggers de novo alveologenesis in a BPD model: impact on the resident mesenchymal niche cells

**DOI:** 10.1101/2022.03.14.484213

**Authors:** Sara Taghizadeh, Cho-Ming Chao, Stefan Guenther, Lea Glaser, Luisa Gersmann, Gabriela Michel, Simone Kraut, Kerstin Goth, Janine Koepke, Monika Heiner, Ana Ivonne Vazquez-Armendariz, Susanne Herold, Christos Samakovolis, Norbert Weissmann, Francesca Ricci, Giorgio Aquila, Laurent Boyer, Harald Ehrhardt, Parviz Minoo, Saverio Bellusci, Stefano Rivetti

## Abstract

Bronchopulmonary dysplasia (BPD) is a neonatal lung disease developing in premature babies characterized by arrested alveologenesis and associated with decreased Fibroblast growth factor 10 (FGF10) expression. One-week hyperoxia (HYX) exposure of newborn mice leads to a permanent arrest in alveologenesis. To test the role of Fgf10 signaling to promote de novo alveologenesis following hyperoxia, we used transgenic mice allowing inducible expression of *Fgf10* and recombinant FGF10 (rFGF10) protein delivered intraperitoneally. We carried out morphometry analysis, and IF on day 45. Alveolospheres assays were performed co-culturing AT2s from normoxia (NOX) with FACS-isolated Sca1^Pos^ resident mesenchymal cells (rMC) from animals exposed to NOX, HYX+PBS, or HYX+FGF10. scRNAseq between rMC-Sca1^Pos^ isolated from NOX and HYX+PBS were also carried out. Transgenic overexpression of Fgf10 and rFGF10 administration rescued the alveologenesis defects following HYX. Alveolosphere assays indicate that the activity of rMC-Sca1^Pos^ is negatively impacted by HYX and partially rescued by rFGF10 treatment. Analysis by IF demonstrates a significant impact of rFGF10 on the activity of resident mesenchymal cells. scRNAseq results identified clusters expressing *Fgf10, Fgf7, Pdgfra*, and *Axin2*, which could represent the rMC niche cells for the AT2 stem cells. In conclusion, we demonstrate that rFGF10 administration is able to induce de-novo alveologenesis in a BPD mouse model and identified subpopulations of rMC-Sca1^Pos^ niche cells potentially representing its cellular target.

## 1 Introduction

Bronchopulmonary dysplasia (BPD) is a complex chronic respiratory disease in preterm infants with no curative treatment and is associated with a high morbidity-mortality rate. Invasive mechanical ventilation, oxygen toxicity, infections, and inflammation are known risk factors for BPD pathogenesis [1]. While the lung of normal infants progresses from the saccular stage to the alveolar stage leading to the formation of the alveoli, the lungs of infants suffering from BPD are characterized by arrested lung development at the saccular stage resulting in immature lungs displaying an emphysematous-like appearance upon histological examination [2].

The cellular mechanisms at play during alveologenesis are currently unclear. Still, it is proposed that the alveolar epithelial type 2 cells (AT2s) could take center stage in mediating the required interactions with the surrounding vascular system and the resident mesenchymal cells to allow the transition from the saccular to the alveolar stage [3]. AT2s play a critical role for the maintenance of efficient gas exchange during homeostasis and serve as progenitor cells for the re-epithelialization of the alveoli after lung injury. In addition, AT2s modify the inflammatory response by secreting different growth factors and cytokine [4].

Fibroblast growth factor 10 is an essential factor expressed by the lung mesenchyme regulating the branching process of the embryonic lung [5]. It acts via the tyrosine kinase receptor Fgfr2b. It has been reported to target both the alveolar epithelial progenitor cells to control their proliferation and differentiation [6] and resident lung mesenchymal cells (rMC) to control their differentiation along the lipofibroblast lineage [7]. Fgf10 expressing lipid-droplet rich resident mesenchymal cells located close to the AT2s are defined as lipofibroblasts and have been proposed to represent the niche for AT2 cells [7]. Indeed, using the alveolosphere organoid model (AT2-rMC co-culture in Matrigel), we have identified rMC-Sca1^Pos^Fgf10^Pos^ cells out of the rMCs to be responsible for the maintenance of AT2 stem cell proliferation and differentiation [8].

Multiple studies have shown that Fgf10 plays a vital role in lung disease and regeneration after injury. For example, in the mouse model of bleomycin-induced lung fibrosis, *Fgf10* overexpression during fibrosis formation or fibrosis resolution demonstrated a protective and therapeutic effect, respectively [9]. Fgf10 is also involved in the regeneration of the bronchial lung epithelium after naphthalene injury [10,11]. Suggestive of a causative role in the bronchial airway manifestations in the context of chronic obstructive pulmonary disease, patients with heterozygous loss of function of *FGF10* exhibit a significant decrease in IVC, FEV1, and FEV1/IVC quota compared to non-carrier siblings and predicted reference values [12].

Supporting this observation, an Fgf10-Hippo epithelial-mesenchymal crosstalk maintains and recruits lung basal stem cells in the conducting airways [13]. While transient *Fgf10* expression by ASMCs is critical for proper airway epithelial regeneration in response to injury, sustained Fgf10 secretion by the ASMC niche, in response to chronic Ilk/Hippo inactivation, results in pathological changes in airway architecture [13]. We also recently identified a new population of repair supportive mesenchymal cells (RSMCs), distinct from the ASMCs but which also secrete Fgf10 to repair the bronchial epithelium [11]. Therefore, Fgf10/Fgfr2b signaling is an exciting target for treating lung pathologies involving the bronchial epithelium.

Furthermore, we experimentally demonstrated in mice that decreased Fgf10 expression is causative for the lethality of newborn mice observed following hyperoxia exposure [14]. Hyperoxia (HYX) exposure (85% O_2_) to newborn mice for 8 days is used to trigger arrested lung development, thereby mimicking the bronchopulmonary dysplasia (BPD) phenotype. Inhibition of Fgfr2b signaling postnatally in the context of HYX also leads to increased alveolar defects and a pulmonary hypertension phenotype [2].

In humans, the decrease in the expression of FGF10 protein in the lungs of patients suffering from BPD suggests a role for FGF10 in the etiology of this disease.[15] FGF10 expression has been reported to be a target of the inflammatory process in BPD [16]. Interactions between NF-κB, Sp1, and Sp3 led to inhibition of *Fgf10* expression [17]. *Fgf10* inhibition is mediated by toll-like receptor 2 and 4 (Tlr2 or Tlr4) activation [15]. Therefore, we hypothesize that the exogenous application of recombinant FGF10 (rFGF10) could potentially represent a solid therapeutic approach. In the present study, we used two in vivo models to test this hypothesis. First, we used transgenic lines to exert gain of function of Fgf10 signaling following hyperoxia injury in newborn mice. Second, we administered rFGF10 protein intraperitoneally or PBS (control) at three different time intervals (P8-P14, P22-P28, and P36-P42) in mice following HYX. We carried out morphometry analysis, and IF to assess the impact of the injury on day 45. We took advantage of 3D organoid cell culture to test the effect of HYX injury on rMC-Sca1^Pos^ cells functionally.

Single cell RNA sequencing (scRNAseq) by using FACS-sorted rMC-Sca1^Pos^ cells at P45 from the NOX or HYX-PBS lungs was carried out and allowed to define potential subpopulations of the Sca1^Pos^ resident mesenchymal cells representing the niche for alveolar type 2 (AT2) stem cells.

## 2 Materials and Methods

### 2.1 Experimental animals

Pregnant C57BL/6J at embryonic day (E) 14 were purchased from Jackson through Charles Rivers Laboratory, Germany. Mice were housed by the Animal Care and Veterinary Service of the University of Justus-Liebig University in accordance with institutional guidelines. Newborn mouse pups from dams that were delivered on the same day were randomized at the day of birth [postnatal day P0] and divided into equal-sized litters of 6 to 8 pups. Following randomization, mice cages were either maintained in room air (normoxia, NOX, 21% O_2_) or in normobaric hyperoxia (HYX, 85% O_2_) from P0 until P8. The hyperoxic environment was maintained in a sealed plexiglass A-chamber with continuous oxygen monitoring (BioSpherix Ltd, Parish, NY). Nursing dams were rotated between NOX and HYX groups every 24 hours to avoid oxygen toxicity and associated confounding factors. All mice were maintained in 12/12 h light/dark cycle and received food ad libidum. At the end of the HYX exposure (P8), pups in the HYX group were subsequently divided into HYX-PBS and HYX-FGF10. Pups in HYX-FGF10 group received 6μg (60 μl) intraperitoneal (i.p.) injection of hrFGF10 (Cat Nr # 345-FG, R&D systems, USA) at P8, P10, P12, P14 followed by having one-week rest. At P22, P24, P26, P28, and P36, P38, P40, P42, mice received 10 μg (100 μl) of hrFGF10. The rationale for the regular time intervals and dosage was based on previous studies [18–20]. Then at P45, all developing mice were sacrificed using CO_2_ and cervical dislocation. All animal studies were performed according to protocols approved by the Animal Ethics Committee of the Regierungspraesidium Giessen (permit numbers: G29/2020-No. 994_GP and G85/2019-No. 986_GP).

*Rosa26*^*rtTA*^*/+;Tg(tet(O)Fgf10)/+* and *Rosa26*^*rtTA*^*/+;Tg(tet(O)sFgfr2b)/+* were obtained from Dr. J.Whitsett (Cincinnati Children’s Hospital Medical Center, Cincinnati, OH, USA), and the experiment were carried out at Childrens Hospital Los Angeles (Licence number 253-08).

### 2.2 Alveolar morphometry

For alveolar morphometry, the blood from the lungs was flushed via the heart first with cold PBS and then with 4% paraformaldehyde (PFA) in phosphate-buffered saline (pH 7.0) at a vascular pressure of 20 cm H_2_O. PBS followed by 4% PFA was infused into the lung via the trachea at a pressure of 20 cm H_2_O. Investigations were performed using 5 μm sections of paraffin-embedded left lobe of the lungs. The mean linear intercept, mean air space, and mean septal wall thickness were measured after staining with hematoxylin and eosin (H&E).

Total scans from the left lobe were analyzed using a Leica DM6000B microscope with an automated stage according to the procedure previously described (McGrath-Morrow et al., 2004; Woyda et al., 2009) which was implemented into the Qwin V3 software (Leica, Wetzlar, Germany). Horizontal and vertical lines (distance 40 μm) were placed across each lung section. The mean linear intercept corresponds to the total length of the horizontal (or vertical) lines present in the section divided by the number of times these lines cross a septal wall. Bronchi and vessels above 50 μm in diameter were excluded prior to the computerized measurement. The air space was determined as the non-parenchymatous non-stained area. The septal wall thickness was measured as the length of the line perpendicularly crossing a septum. From the respective measurements, mean values were calculated.

### 2.3 High-resolution Echocardiography

Transthoracic echocardiography was performed with a Vevo770 high-resolution imaging system to measure right ventricular wall thickness (RVWT), right ventricular internal diameter (RVID), tricuspid annular plane systolic excursion, and pulmonary artery acceleration time. At day 45, transthoracic echocardiography was performed under anaesthesia, induced with 3% isoflurane and maintained via a nose cone with 1.5% isoflurane (balanced with O2).

### 2.4 Lung dissociation and fluorescence-activated cell sorting

Lungs from mice at P45 were collected and processed for single cell suspension. Samples were prepared for sorting by FACSAria III (BD Bioscience) based on an established protocol previously described in detail [8].

### 2.5 Alveolar Organoid assay

rMC-Sca1 mesenchymal sorted cells from male mice of different groups; NOX, HYX+PBS, and HYX+FGF10 were combined with AT2 mature from NOX group. 5000 Cd45^Neg^Cd31^Neg^Epcam^Pos^Lyso^Pos^cells and 50000 Cd45^Neg^Cd31^Neg^Epcam^Neg^Sca1^Pos^ cells were resuspended in 100 μl culture medium (sorting media plus 1% ITS (gibco # 41400-045)) and mixed 1:1 with 100 μl growth factor-reduced phenol Red-free Matrigel (Corning #356231). Cells were seeded in individual 24-well 0.4 μm Transwell inserts (Falcon, SARSTEDT). After incubation at 37°C for 15 minutes, 500 μl of culture was placed in the lower chamber and the plate was placed at 37°C in 5% CO_2_/air.

The culture medium was changed every other day. ROCK inhibitor (10 μM, Y27632 STEMCELL#72304) was included in the culture medium for the first two days of culture. Organoids were counted and measured at day 14.

### 2.6 Whole-mount immunofluorescence staining of organoids

Organoids (at day 14) were fixed in 4% paraformaldehyde for 30 min followed by 3x washing steps with PBS and incubation in 0.1% Triton X-100 for 30 min. After washing 3x with PBST, organoids were blocked with 1xTBS, 3% BSA. 0.4% Triton X-100 for 1 hr at room temperature. Organoids were washed and then incubated at 4°C overnight with 1x TBS, 1.5% BSA, 0.2% Triton X-100, and primary antibody against Sftpc (AB3786, Millipore; 1:500), Pdpn (8.1.1, DSHB; 1:250). The next day, after 3x washing with TBST for 10 min, organoids were incubated with secondary antibodies (AlexaFlour 488 goat anti-Rabbit IgG Green (1:500) Cat. #11034 invitrogen) at RT and washed three times with TBST before being mounted with Prolong Diamond Anti-fade Mountant with DAPI (Invitrogen 4’,6-diamidino2-phenylindole). Photomicrographs of immunofluorescence staining were taken using a Leica DMRA fluorescence microscope with a Leica DFC360 FX camera (Leica, Wetzlar, Germany). Figures were assembled using Adobe Illustrator.

### 2.7 Immunofluorescence staining

For immunofluorescence staining, the slides were deparaffinized, blocked with 3% bovine serum albumin (BSA) and 0.4% Triton X-100 in Tris-bufered saline (TBS) at room temperature (RT) for 1 h and then incubated with primary antibodies against Sftpc (AB3786, Millipore; 1:500), Pdpn (8.1.1, DSHB; 1:250) at 4 °C overnight. After incubation with primary antibodies, slides were washed three times in TBST (Tris buffer saline+0.1% Tween 20) for 5 min, incubated with secondary antibodies at RT and washed three times with TBST before being mounted with Prolong Diamond Anti-fade Mountant with DAPI (4′,6-diamidino-2-phenylindole; Invitrogen).

### 2.8 scRNA-seq library preparation

Single-cell suspensions were processed using the 10x Genomics Single Cell 3′ v3 RNA-seq kit. Gene expression libraries were prepared according to the manufacturer’s protocol. MULTI-seq barcode libraries were retrieved from the samples, and libraries were prepared independently.

### 2.9 Sequencing and processing of raw sequencing reads

Sequencing was done on Nextseq2000, and raw reads were aligned against the mouse genome (mm10, ensemble assembly 104) and mapped and counted by StarSolo (Dobin et al., doi: 10.1093/bioinformatics/bts635) followed by secondary analysis in Annotated Data Format. Pre-processed counts were analyzed using Scanpy (Wolf et al., doi: 10.1186/s<u>13059-017-1382</u>-0). Primary cell quality control was conducted by taking the number of detected genes and mitochondrial content into consideration. We removed 232 cells in total that did not express more than 300 genes or had a mitochondrial content of less than 20%. Further, we filtered genes if detected in less than 30 cells (<3%). Raw counts per cell were normalized to the median count over all cells and transformed into log space to stabilize the variance. We initially reduced the dimensionality of the dataset using PCA, retaining 50 principal components. Subsequent steps, like low-dimensional UMAP embedding (McInnes & Healy, https://arxiv.org/abs/1802.03426) and cell clustering via community detection (Traag et al., https://arxiv.org/abs/1810.08473), were based on the initial PCA. Final data visualization was done by cellxgene package.

### 2.10 Statistical analysis

Data are presented as means ± SD. All statistical analyses were performed with GraphPad Prism 8.0. ROUT method for outliers’ identification was performed for each experimental group. Comparison between three experimental groups was made using one-way ANOVA analysis with multiple comparisons between three groups.

Comparison between HYX+PBS and HYX+FGF10 was done using Student’s t-test.

*P* values < 0.05 were considered as significant and depicted as following: *P* values < 0.05: *; *p* values < 0.01: **; *p* values < 0.001: ***; *p* values <0.0001: ****.

## 3 Results

### 3.1 Overexpression of Fgf10 in vivo promotes de novo alveologenesis after hyperoxia injury in newborn pups

First, we validated in our experimental conditions the consequences of hyperoxia (HYX) exposure (85% O_2_) from postnatal day 2 (P2) to P9 of newborn WT C57BL6 mice on postnatal lung development. Newborn animals exposed to normoxia (NOX) were used as controls. Consequently, animals from the NOX and HYX groups at P9 were kept in NOX conditions up to P45 (Fig. 1A). Then, the animals were sacrificed at P9, P15, and P45, and lung morphometry analysis was carried out (Fig. 1B,C). The mean linear intercept (MLI) quantification shows a significantly higher value in HYX vs. NOX lungs at P9 (n=4 animals per time point in each group were used 21.0 μm vs. 28.1 μm, *p*=0.013). This increase in MLI persists at P15 and P45 (17.6 μm vs. 23.2 μm, *p*=0.011; 14.6 μm vs. 26.1 μm, *p*=0.016) when the previously exposed HYX animals are kept in NOX. Consistent with the increase in MLI, we also found a trend towards an increase in the airspace at P9 and P15 and a significant increase at P45 (41.4% vs. 59.7%, *p*=0.041). No significant differences in the septal wall thickness were observed in this experiment. In conclusion, HYX exposure leads to arrested lung development. Furthermore, the permanence of the defects triggered by HYX is similar to what is observed in the lungs of infants with BPD.

**Figure 1.**
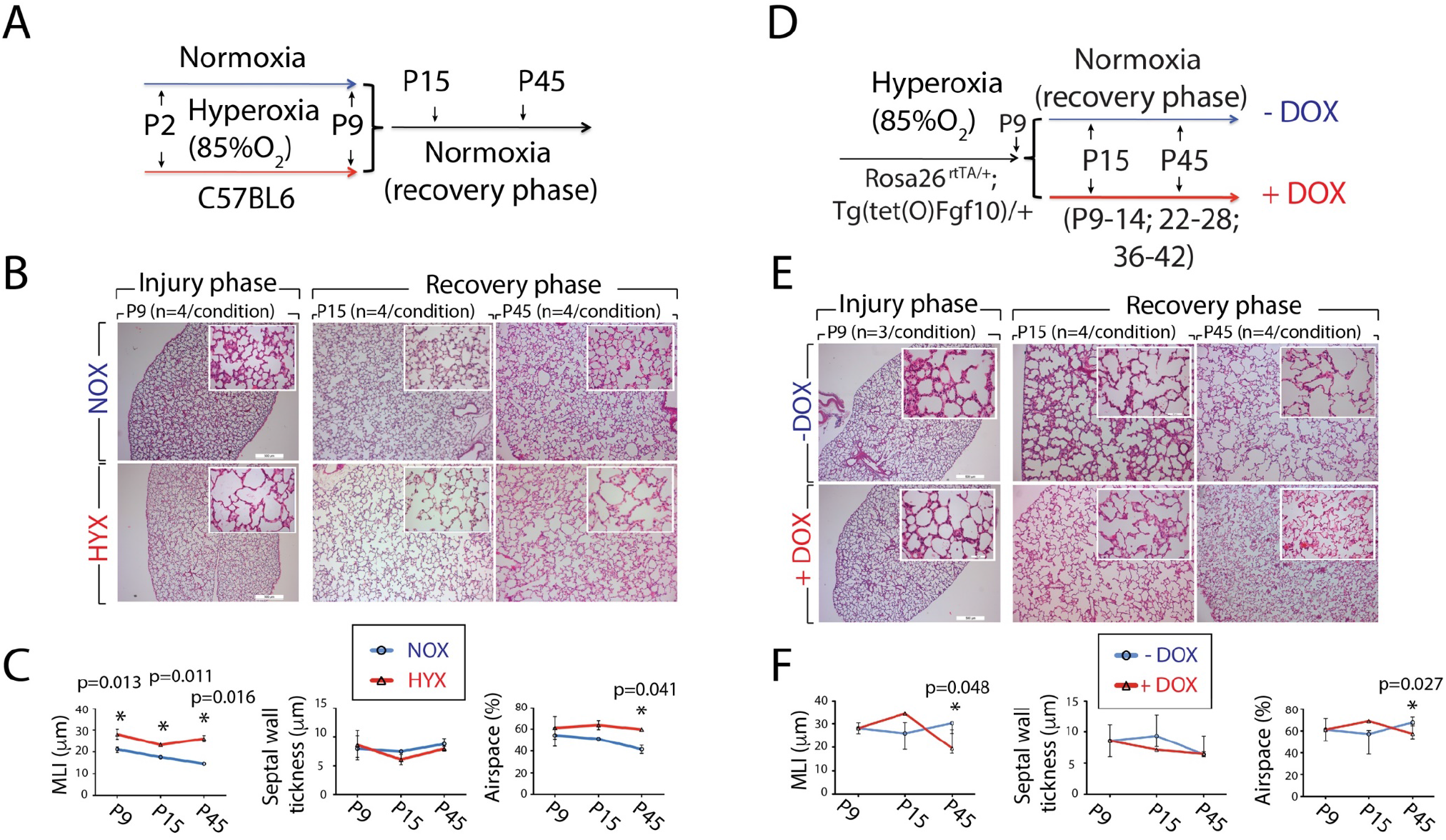

Next, we used a double transgenic line (*Rosa26*^*rtTA*^*/+;Tg(tet(O)Fgf10)/+*) to ubiquitously express *Fgf10* (Fig. 1D-F). This line was previously validated in the lung in the context of fetal development and the adult mouse [9,10]. *Rosa26*^*rtTA*^*/+;Tg(tet(O)Fgf10)/+* mice were exposed to HYX from P2 to P9 and subsequently divided into two groups put in NOX conditions for up to P45. During this “recovery” period, the experimental group was exposed to DOX (through the mother’s milk as the mother is fed with doxycyclin food), while the lactating females of the control group were exposed to regular food. The design of this experiment allowed us to evaluate the impact of Fgf10 overexpression on inducing de novo alveologenesis. Morphometric quantification at P45 between experimental (+DOX) vs. control (-DOX) groups (n=4 animals per time point and condition) demonstrated a significant decrease in the MLI (30.5 μm vs. 21.4 μm, *p*=0.048) and airspace (68.3% vs. 58.5%, *p*=0.027) with no substantial change in the septal wall thickness.

These results are in harmony with the observed increased alveolarization in the experimental vs. control group during the “recovery” period. Altogether, these results support the use of FGF10 to induce de novo alveologenesis in the context of experimental BPD.

### 3.2 Fgfr2b signaling is not involved in the endogenous repair process after hyperoxia injury

The permanence of the defects triggered by HYX when the newborn mice are re-exposed to NOX suggests that “repair” process per se is inefficient. However, we cannot exclude that such a “repair” process is involved in maintaining a steady-state situation. Therefore, the expectation is that inhibiting this repair process will worsen the lung structure.

We previously reported that Fgfr2b ligands (*Fgf1, 3, 7*, and *10*) are expressed in the lung between P0 and P3 [2]. Upon HYX exposure, these ligands are impacted (with an increase in *Fgf3* and *Fgf1* and a decrease in *Fgf10* during the P3-P8 HYX exposure period). To determine if Fgfr2b ligands were still functional after HYX injury to maintain the steady-state (“repair”) of the lung during the “recovery” period (NOX exposure), we used a dominant-negative transgenic mouse model; *Rosa26*^*rtTA*^*/+;Tg(tet(O)sFgfr2b)/+*. This loss of function approach exploits the features of a dominant-negative soluble form of Fgfr2b (sFgfr2b), which expression is induced by DOX. sFgfr2b traps all Fgfr2b ligands, preventing them from acting on their endogenous receptors.

Following HYX exposure, animals are re-exposed to NOX. From P9 onwards, the experimental group was exposed to DOX while the control group was exposed to regular food (Fig 2A), and the corresponding lungs were examined at P9, P15, and P45 by morphometry (Fig 2B-C). No significant difference in MLI, septal wall thickness, and airspace level were observed between the two groups, indicating that endogenous Fgfr2b signaling does not maintain the steady-state following HYX.

**Figure 2.**
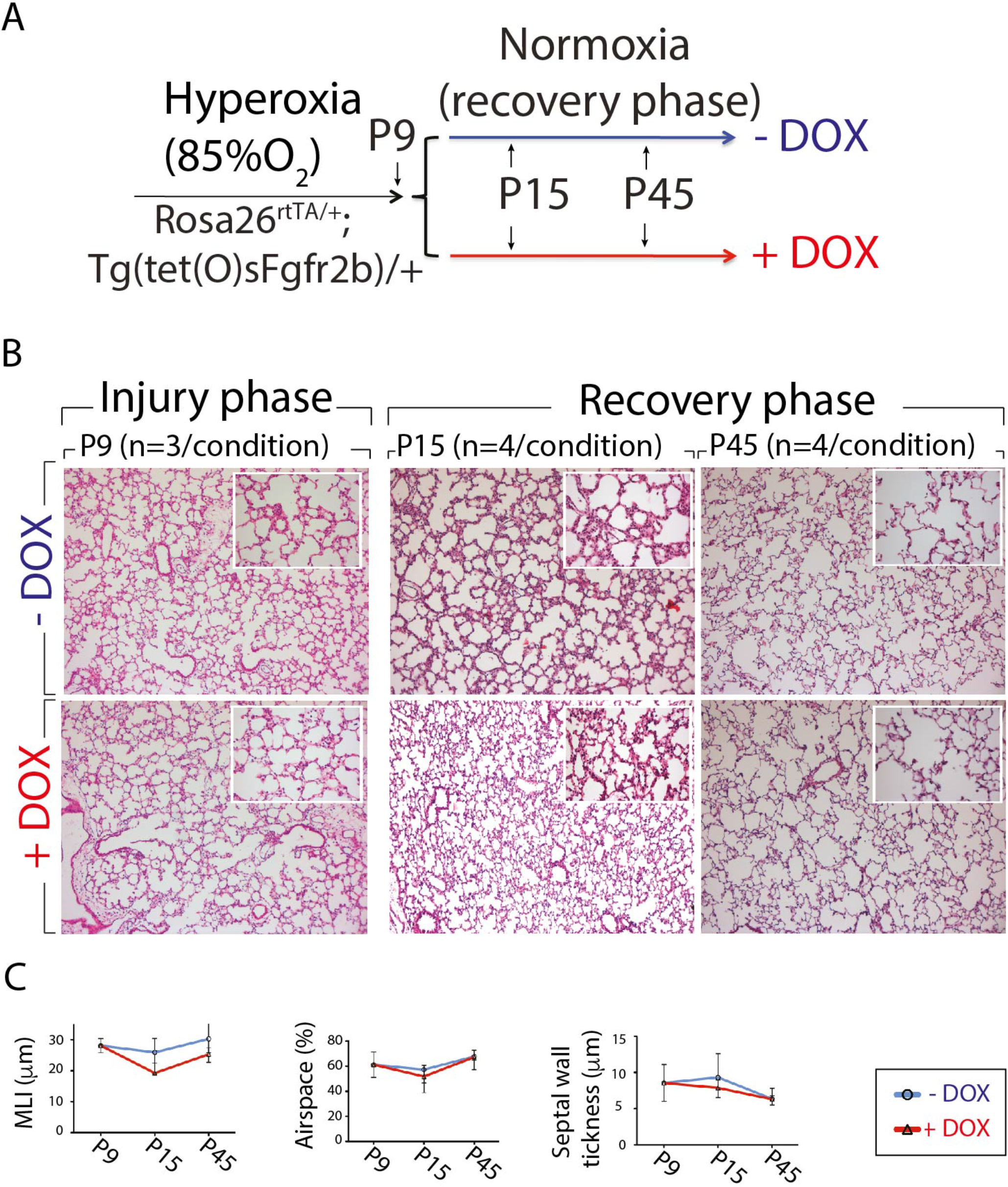

### 3.3 Human recombinant FGF10 rescues the lung structure in mice following hyperoxia induced-arrest in alveologenesis

To test the therapeutic potential of human recombinant FGF10 (rFGF10) in the repair process after HYX injury, we delivered rFGF10 to wild-type mice. Figure 3A shows the experimental scheme depicting the injury period and time points of rFGF10 administrations.

**Figure 3.**
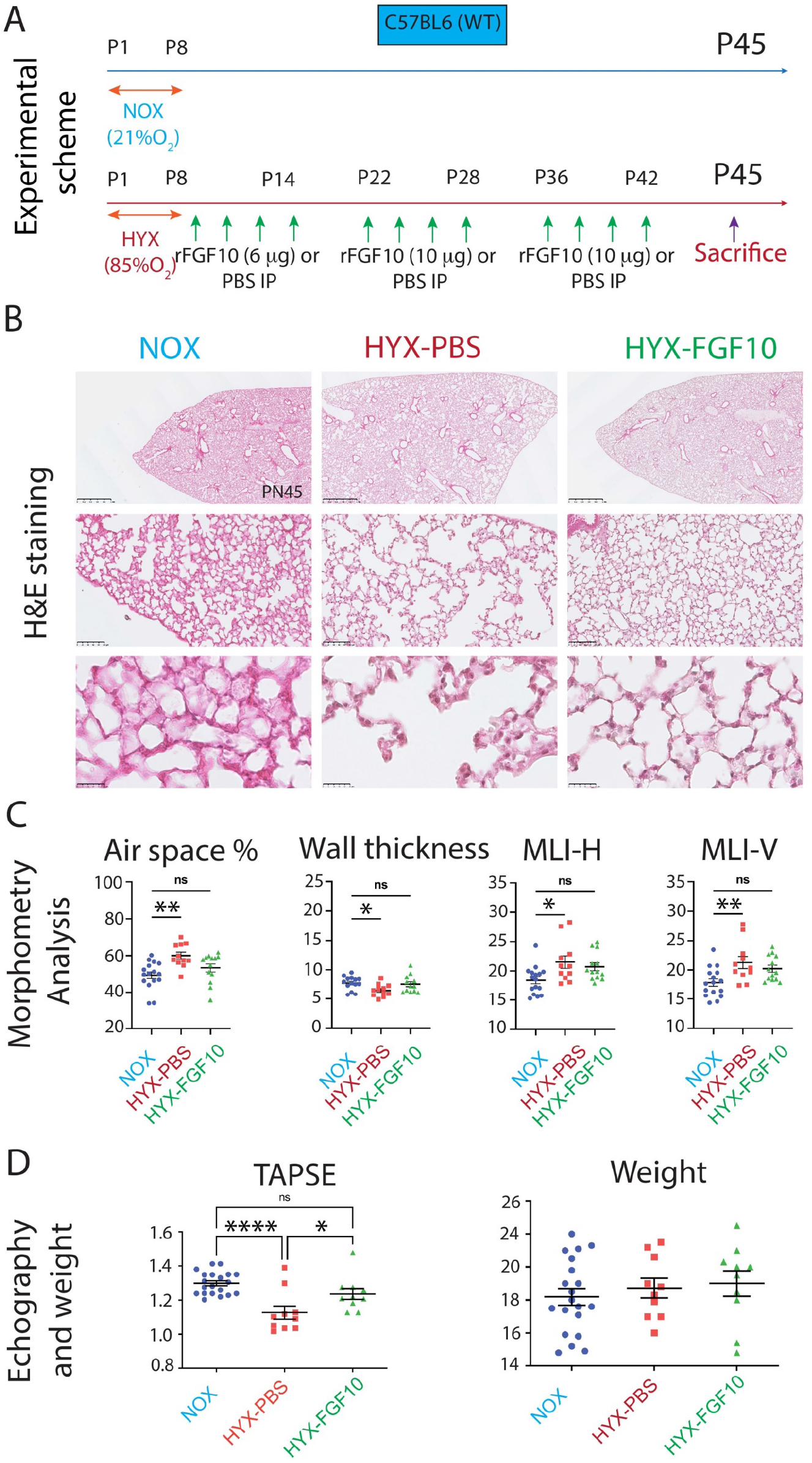

C57BL6 pups from the experimental group were exposed to HYX (85% O_2_) from P0 to P8. In parallel, newborn mice belonging to the control group were maintained under room air (21% O_2_). At P8, pups from HYX experimental group were re-exposed to NOX and subsequently divided into two groups, either receiving PBS (HYX+PBS) or rhFGF10 (HYX+FGF10) via intraperitoneal injections (i.p) at three different intervals every other day (P8-P14, P22-P28, and P36-P42). The dose of rFGF10 was 6 μg/g at the first period and 10 μg /g for the second and third treatment periods. The timing of administration and the dosage of rFGF10 were based on previous studies [18–20].Mice were sacrificed at P45. In addition, echocardiography to assess heart function as a surrogate for pulmonary arterial hypertension and morphometry analysis to assess architectural defects were carried out.

Morphometric analysis of H&E sections (from NOX (n=16), HYX+PBS (n=11) and HYX+FGF10 (n=13) groups was carried out. Our results indicate arrested lung development between the HYX+PBS and NOX lungs. Increased airspace (p=0.0021), decreased wall thickness (p=0.048), and increased MLI (both horizontal and vertical) (p=0.0175, p=0.0066) are observed in HYX+PBS vs. NOX lungs (Fig 3B, C) indicating that hyperoxia led to the expected damages. Supporting the role of FGF10 in de-novo alveologenesis (suggested in Fig. 1D-F), an obvious recovery of the lung structure was observed in HYX+FGF10 vs. HYX+PBS lungs (Fig 3B). Interestingly, ANOVA analysis did not indicate significant changes for morphometric parameters between these two groups (Fig 3C). However, a direct comparison between the HYX+PBS and HYX+FGF10 group by student t-test showed a statistically significant difference in air space (p=0.03), as well as a trend towards significance for wall thickness (*p*=0.064). However, no significance was noted for MLI-H (*p*=0.522) and MLI-V (*p*=0.369) (Supp-Fig 1).

BPD is correlated with abnormalities in the right ventricle (RV). These heart defects are usually associated with PAH formation. RV function was evaluated by using echocardiography. In particular, we measured the tricuspid annular plane systolic excursion (TAPSE, n= NOX:21, HYX+PBS:11, HYX+FGF10:11) value <1.6 mm, as this was reported to be a sensitive method to estimate RV systolic dysfunction. In HYX+PBS vs. NOX, we observe a significant decrease in TAPSE (*p*<0.00001). As the TAPSE value is highly correlated with the body weight, we also monitored the weight of the pups between the two groups. No significant difference was found between them (Fig 3D). Comparison between the HYX+FGF10 vs. the HYX+PBS groups indicates an increase of the TAPSE value (p=0.028).

No significant difference was found between the HYX+FGF10 vs. the NOX groups (*p*=0.1925), indicating a normalization of the RV function. This difference between the HYX+FGF10 and HYX+PBS groups was not due to changes in body weight (*p*=0.959). Taken together, these observations support the therapeutic potential of rFGF10 delivered IP in the *de novo* alveologenesis process after hyperoxia injury.

### 3.4 Alveolospheres assays indicate that HYX and FGF10 pre-treatments impact FACS-isolated rMC-Sca1^Pos^ activity to sustain AT2 stem cell proliferation and differentiation

Alveolar epithelial type 2 cells (AT2) are the quiescent stem/progenitor epithelial cells capable of proliferating and differentiating to alveolar epithelial type 1 (AT1) cells following exposure to injury. Resident mesenchymal cells (rMC) have been previously defined as Cd45^Neg^Cd31^Neg^Epcam^Neg^ population and represent at large a niche for AT2 stem cells. In particular, rMC-Sca1^Pos^ cells effectively trigger organoid formation using the alveolosphere assay [8]. To perform this assay, we have collected lungs at P45 from NOX, HYX+PBS, and HYX+FGF10 mice and sorted the Cd45^Neg^Cd31^Neg^Epcam^Neg^ Sca1^Pos^ cells (rMC-Sca1^Pos^). The rMC-Sca1^Pos^ cells belonging to these three groups were individually co-cultured in Matrigel with Epcam^Pos^Lyso^Pos^ cells from wild type, non-injured lungs representing mature AT2 cells and analyzed after 14 days in culture (Fig 4A).

**Figure 4.**
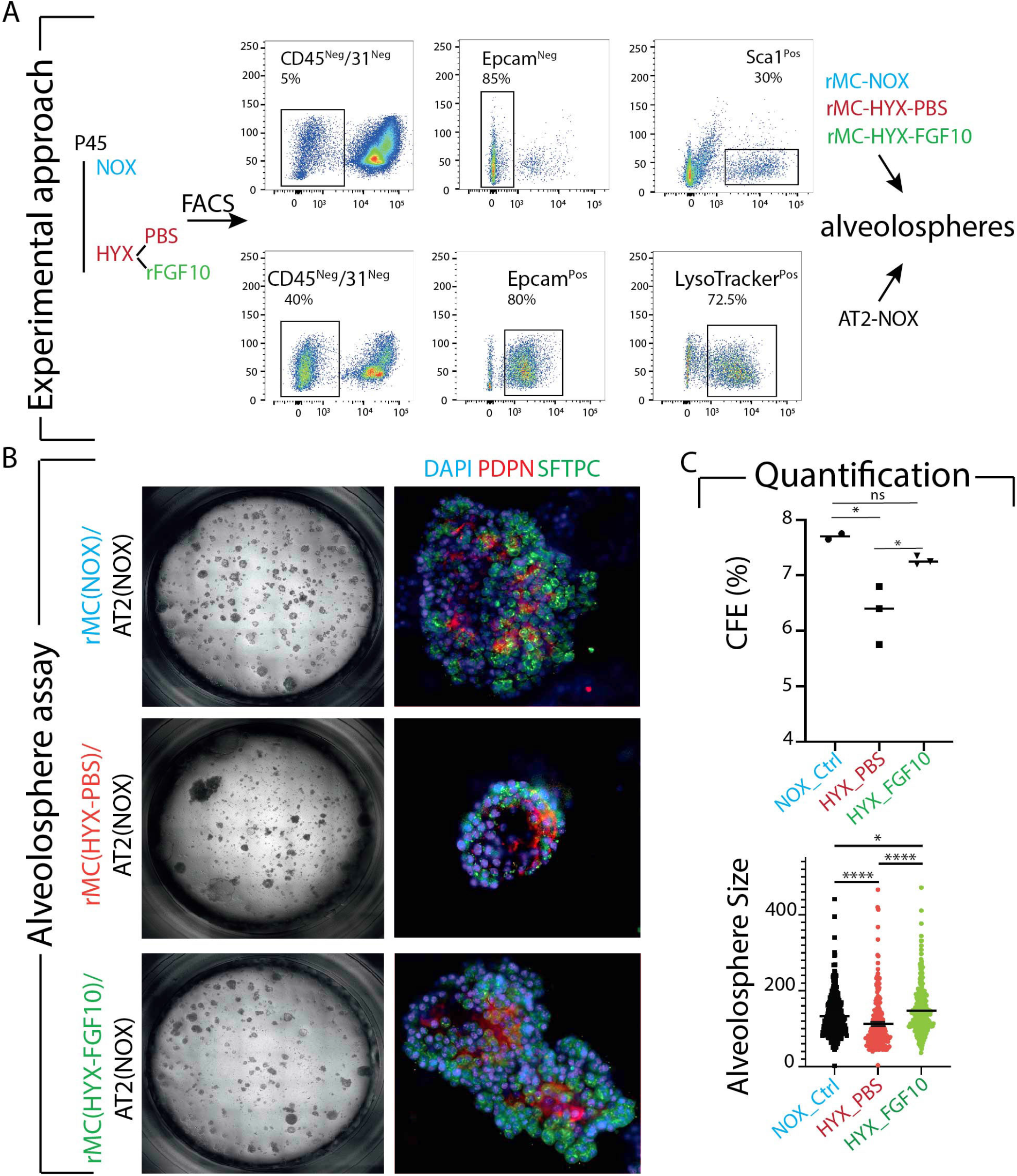

Colony forming efficiency (CFE), which corresponds to the ratio of the number of spheres that grow out of the total number of seeded epithelial progenitor cells as well as organoid size were measured for each group. Comparison between rMC-Sca1^Pos^ (HYX+PBS) and rMC-Sca1^Pos^ (NOX) indicates a decrease in both CFE (n=3, *p*< 0.0001) and organoid size (n=3, *p*< 0.0001). Comparison between rMC-Sca1^Pos^ (HYX+FGF10) and rMC-Sca1^Pos^ (HYX+PBS) indicates a significant increase both in organoid formation (CFE) (n=3, *p*< 0.01) and in organoid size (n=3, *p*< 0.0001).

Next, we quantified by IF the number of Pdpn^Pos^ (AT1 cells) and Sftpc^Pos^ (AT2 cells) out of total cells in organoids in our three conditions. Examination of Pdpn^Pos^ cells indicates that no significant difference was found between the HYX+PBS vs. the NOX groups. However, comparison between the HYX+FGF10 vs. the HYX+PBS groups indicates a decrease of Pdpn^Pos^ cells in the HYX+FGF10 group (*p*=0.0001). Similarly, analysis of Sftpc^Pos^ cells shows that there is no significant difference between the HYX+PBS vs. the NOX groups. By contrast, there is significant increase of Sftpc^Pos^ cells in the HYX+FGF10 groups compared to both the HYX+PBS and NOX groups (*p*=0.0007) (Figure 4 B and C).

In conclusion, the activity of the rMC-Sca1^Pos^ niche in terms of CFE and organoid size is negatively impacted by HYX, and human rFGF10 treatment restores this activity. In addition, rFGF10 treatment also modified the capacity of the rMC-Sca1^Pos^ niche to direct the AT2 stem cells in the organoids towards the AT2 or AT1 lineage with an increased number of AT2s at the expense of the AT1s.

### 3.5 rMC-Sca1^Pos^ cells are quantitatively impacted by HYX and FGF10

The number of rMC-Sca1^Pos^ cells in NOX, HYX+PBS, and HYX+FGF10 was quantified by flow cytometry analysis. Comparison between HYX+PBS vs. NOX indicates a decrease (12±8 vs. 25±5, *p*<0.001, n=3). Comparison between HYX+FGF10 vs. HYX+PBS shows an increase (40 ±4 vs. 12±8, *p*<0.001, n=3). To confirm our results by flow cytometry, we have performed IF staining against Sftpc in the three groups (Fig 5B). Quantification of AT2 cell number in HYX+PBS vs. NOX and HYX+FGF10 vs. HYX+PBS indicated no significant change in total Sftpc^Pos^ cells per total counted cell number (Fig 5C). We also found a considerable decrease of DAPI^Pos^ cells in HYX+PBS group vs. NOX, supporting the alveolar simplification phenotype triggered by HYX. These changes in the number of DAPI^Pos^ cells are partially reversed upon human rFGF10 treatment. Interestingly, we have also found a significant increase in Pdpn^Pos^ cell number in HYX+PBS group versus the NOX group (*p=*0.0002). This increase is rescued by rFGF10 treatment (*p=*0.0001) (Supp-Fig 5). In conclusion, we demonstrated that the mesenchymal compartment represents a significant target of HYX, both qualitatively (Fig 4) and quantitatively (Fig 5).

**Figure 5.**
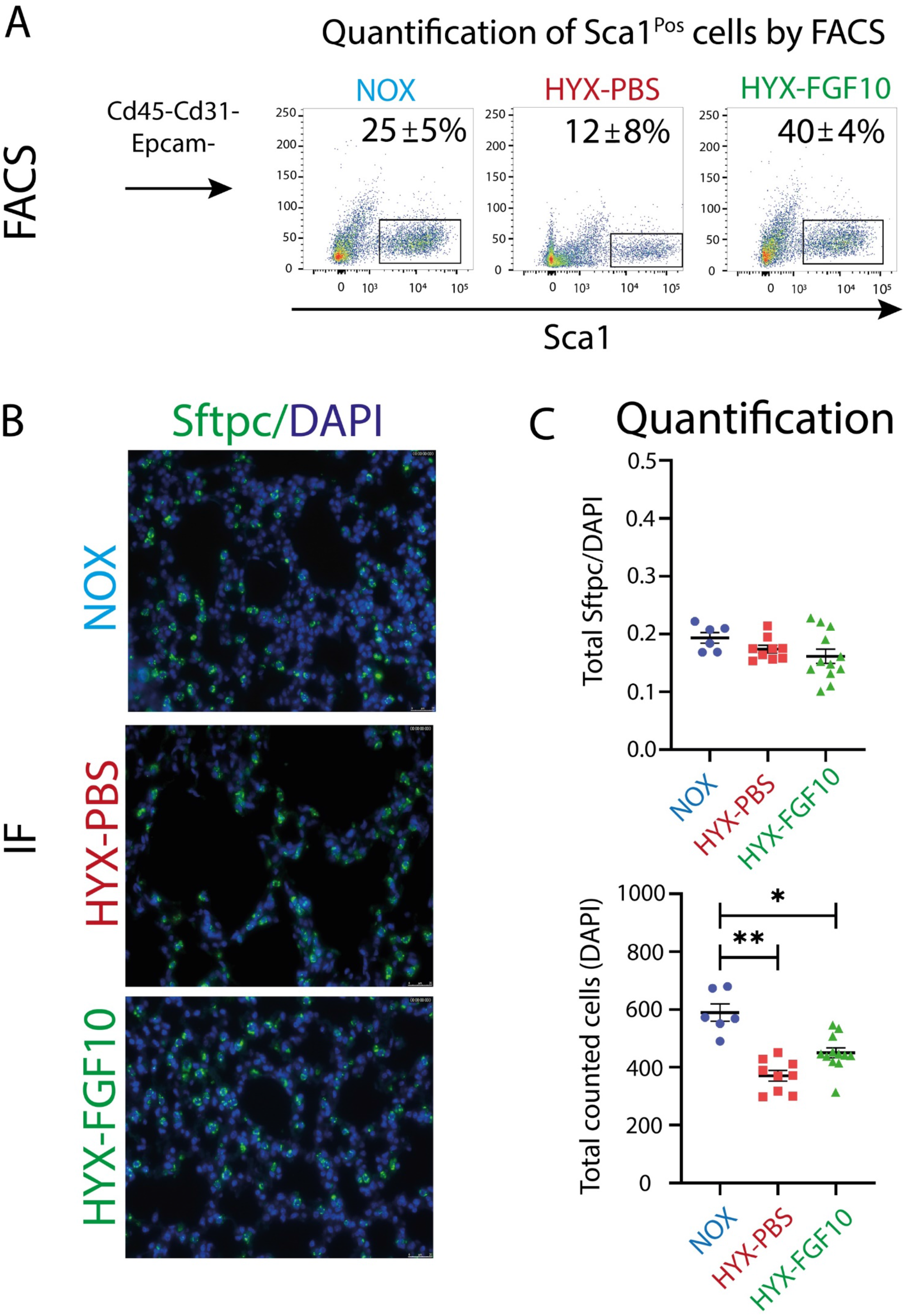

### 3.6 Characterization rMC-Sca1^Pos^ by scRNA-seq in NOX and HYX+PBS

To define different subsets of rMC**-**Sca1^Pos^ cells, we sorted the Cd45^Neg^Cd31^Neg^Epcam^Neg^Sca1^Pos^ cells after HYX+PBS or NOX at P45 and loaded 9000 cells per group on 10x RNAseq chips (Fig 6A). Quality control was performed independently on each library to find the appropriate filtering thresholds. Poor quality cells with a high percentage (>20%) of UMIs mapped to mitochondrial genes were removed. We selected 877 cells in the NOX and 900 in HYX. We detected an average of 2271 and 2389 genes in NOX and HYX with a median number of reads per cell in NOX and HYX of 38.383 and 33.531, respectively (Fig 6B). Performing UMAP embeddings identified a total 18 clusters, which were generated based on initial PCA (Fig 6C). As expected, Ly6a expression was expressed throughout all the different clusters at variable levels, and the epithelial marker Epcam was essentially absent from this data set (Supplementary Fig. 2A).

**Figure 6.**
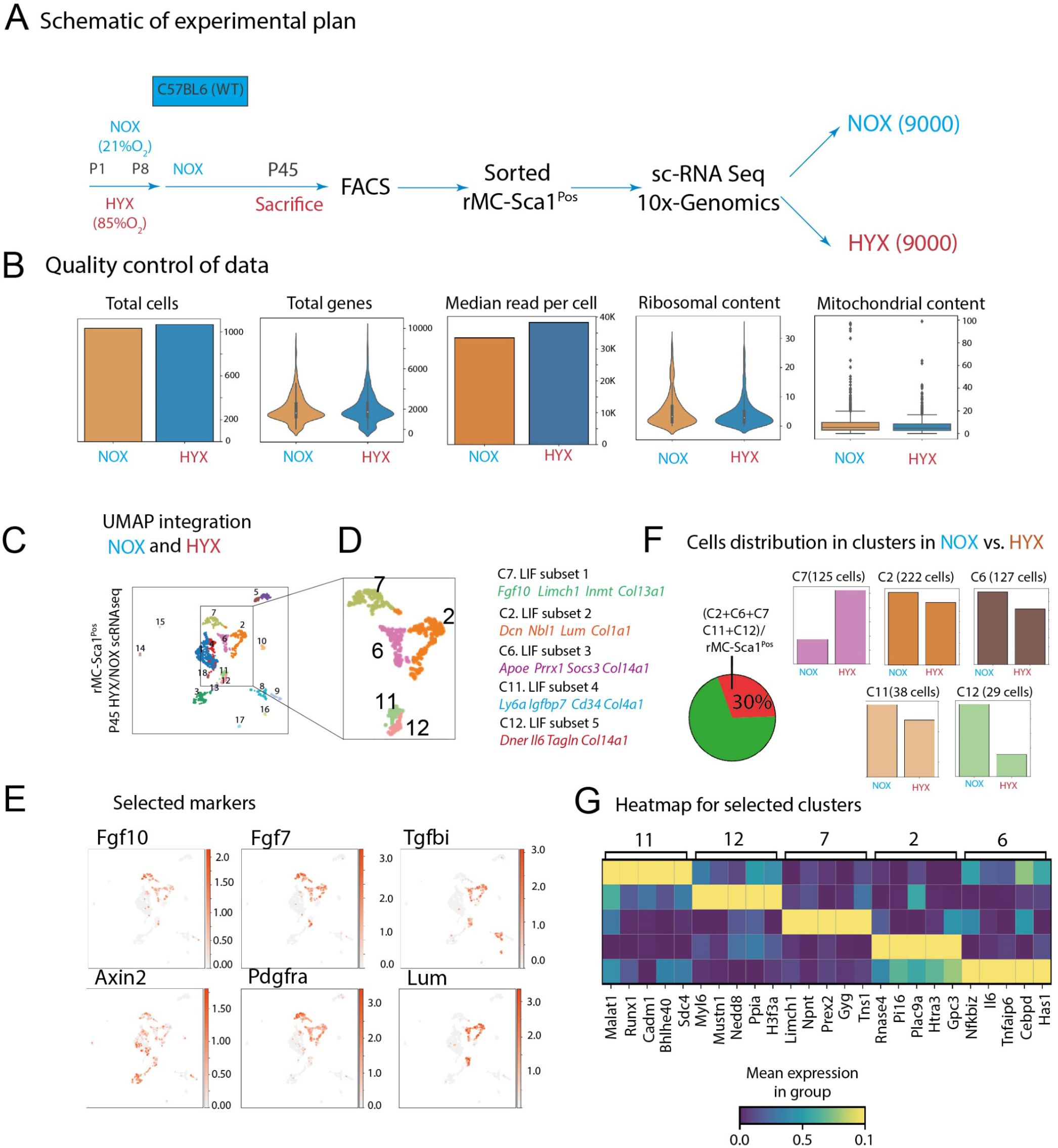

The genes characterizing these other clusters are shown in Supplementary Fig. 2B. These clusters correspond to three major cell groups: lipofibroblasts (LIFs, clusters 2, 6, 7, 11, and 12), myofibroblasts (MYF clusters 1 and 4), and extracellular matrix (ECM) cells (clusters 3 and 13).

Next, we analyzed the expression level of published known markers for the rMC niche [21– 25]. *Fgf10* and *Fgf7* are mainly expressed in the LIF group (clusters 2, 6, and 7). *Tgfbi* and *Lum* are also mainly labeling the LIF group (clusters 2 and 6). *Axin2* is found throughout the rMC-Sca1^Pos^ groups, which indicates that it does not represent a specific marker for the specialized niche cells within this mesenchymal population. *Pdgfra* is mostly expressed in clusters 2, 6, and 7. Finally, *Lumican* is found almost exclusively in clusters 2, 6, and 11-12 (Fig 6E, Supplementary Fig 3). Fig 6G and supplementary Figure 2B display the genes characteristic of each LIF subclusters. Cluster 7 defined as LIF subset 1, significantly expresses *Fgf10, Limch, Inmt*, and *Col13a1*. Cluster 2 as LIF subset 2 expresses *Dcn, Nbl1, Lum*, and *Col1a1* preferentially. Cluster 6 identified as LIF subset 3 is highly enriched for *Apoe, Prrx1, Socs3, Col14a1*. Cluster 11 defined as LIF subset 4 highly expresses *Ly6a* (Sca1), *Igfbp7, Cd34, Col4a1*, Cluster 12 defined as LIF subset 5 is enriched in *Dner, IL6, Tagln* and *Col14a1*.

In our previous study, we found by flow cytometry that around 25% of rMC**-**Sca1^Pos^ cells are Fgf10^Pos^ [8]. rMC**-**Sca1^Pos^ Fgf10^Pos^ cells are the cells which display the rMC niche activity for AT2 stem cells. In our scRNAseq analysis, we found that the LIF group represents 30% of the rMC**-**Sca1^Pos^ cells suggesting that the scRNAseq process conserved the ratio present initially in terms of rMC**-**Sca1^Pos^ Fgf10^Pos^ over rMC**-**Sca1^Pos^ cells.

Among the five clusters in the LIF group, clusters 2 and 6 are the most abundant clusters displaying the highest level of *Fgf10* expression. These clusters show a reduction in cell number after HYX (Fig 6F). An extended list of the genes capable of distinguishing subclusters 2 and 6 from the other LIF subclusters is shown in Supplementary Fig. 2B. Interestingly, cluster 6 is characterized by the expression of genes controlling inflammatory responses like *Cebpd* [26], *Tnfaip6* [27], *Il6* ^20,^ and *Nfkbi* [29].

We then determined the genes differentially expressed in HYX vs. NOX for subclusters 2 and 6 (Supplementary Fig. 4). In general, our analysis indicated that, in our experimental conditions, the range in the expression level for these genes between the two conditions is quite small. Among the genes identified for cluster 2, we noticed *Vegfd*, which is highly expressed during vascular formation and correlates with pulmonary hypertension after hyperoxia injury [30,31]. We also found *Akap12* (A-kinase anchoring protein 12), a gene encoding a protein which function is to bind regulatory subunits of protein kinase A and C and control cells’ growth [32].

Among the genes which were identified for cluster 6, we found proto-oncogene such as *Fosb* and *Jun* [33].

These genes are involved in the regulation of cell proliferation and differentiation. We also found that *Apoe*, a well-accepted LIF marker [22], is expressed more in HYX than NOX. Interestingly, *Nbl1*, a gene encoding an antagonist for BMP signaling [34] is expressed more in NOX than HYX. Altogether, our analysis reveals subtle changes in gene expression in clusters 2 and 6 in HYX vs. NOX, which may be explained by the late time point (P45) chosen for the analysis as well as compensatory mechanisms at play.

## 4 Discussion

So far, there are no therapies for bronchopulmonary dysplasia (BPD), a respiratory disease occurring in preterm infants. As this disease is characterized by a high morbidity and mortality rate, such treatments are urgently needed. Preventive strategies and symptomatic treatments for BPD include innovative ventilation strategies, prenatal corticosteroids, surfactant-replacement, caffeine and vitamin A [35].

The holy grail in the field of lung regeneration is to identify drugs capable of jump-start alveoli formation after injury. In this context, the BPD model, with its permanent arrest in lung development at the saccular stage, is an ideal system to test drugs displaying this capability. Furthermore, the read-out for this de-novo alveologenesis process, primarily based on morphometry analysis, is reliable, easy to perform, and informative. In this context, we have confirmed, using genetic and translational approaches, that FGF10 can induce *de novo* alveologenesis following HYX treatment. This original observation is in agreement with published evidence establishing that 1) FGF10 expression is decreased in lungs of patients with BPD and could therefore be causative for the diseases and that 2) decrease in *Fgf10* expression in mice leads, upon exposure to HYX, to a worsening of the structural defects and lethality.

Fgf10 acts both on the epithelium and mesenchyme [7]. During development, Fgf10 acts on the alveolar progenitor to control their proliferation and differentiation along the alveolar type 2 lineage [6,36,37]. Fgf10 also acts on lung mesenchymal progenitors to control their differentiation towards the lipofibroblast lineage [7]. Suggesting a crucial role for Fgf10 for the control of the proliferation and differentiation of AT2 stem cells in the adult lung, we recently refined the resident mesenchymal cells (defined as Cd45^neg^Cd31^neg^Epcam^neg^) eliciting this activity in vitro using the alveolosphere model. We found that rMC Sca1^Pos^Fgf10^Pos^ cells were the rMC subpopulation displaying most of the activity. These cells were defined as a niche for AT2 stem cells [8]. We also reported that the niche activity of rMC Sca1^Pos^ cells was impacted by metabolic dysfunction, and gender [8].

We initially quantified the activity level of rMC Sca1^Pos^ cells in the context of NOX and HYX+PBS exposure at P45, one month after the HYX exposure. We found a significant drop in the niche activity in the HYX+PBS group, and interestingly, this activity could be partially restored by FGF10 administration. scRNAseq analysis of the rMC Sca1^Pos^ cells in the context of NOX and HYX+PBS allows identifying a subpopulation of the rMC Sca1^Pos^ cells, which co-expressed specifically *Fgf10* and *Fgf7*. Based on the expression of these two markers, we propose that these cells represent the *bona fide* niche for AT2 stem cells.

Interestingly, these cells can be grouped in 2 subclusters (cluster 2 and 6) belonging to the Lipofibroblast cluster. As the analysis was done at P45, the difference in gene expression between the HYX+PBS and NOX groups was subtle. However, we did notice a shrinking in the respective number for these 2 subclusters in HYX+PBS compared to NOX, indicating that HYX could have a quantitative impact at the level of these 2 subclusters. Interestingly, a global effect on the activity of the rMC Sca1^Pos^ cells was observed upon HYX, and this impact was corrected upon FGF10 treatment. In the future, more work will have to be done to better characterize these 2 subclusters at earlier time points following HYX exposure. In addition, a genetic model using dual Dre/Cre recombinase strategies will have to be established to specifically lineage-trace and characterize these 2 subpopulations. In that respect, the *Fgf10*^*CreERT2*^ line that we recently generated and validated will be instrumental in targeting the rMC Sca1^Pos^Fgf10^Pos^ cells [38]. In addition, other Dre lines, with genes expressed specifically in clusters 2 or 6 will have to be generated and validated.

Recently, scRNAseq data on total cells from HYX and NOX lungs at different time points (3, 7, and 14) during HYX exposure (85%O_2_) were reported [25]. Dynamic changes triggered by HYX of the transcriptomic profiles in all the different subpopulations (which included the alveolar epithelium, the resident mesenchymal cells, the endothelium, and the macrophages) were observed. The authors concluded that HYX-induced inflammation was the main force behind these changes. Subsequently, using this dataset, the authors analyzed a minimal number of rMC Sca1^Pos^ cells (which in their paper are named Ly6a positive resident lung mesenchymal stromal cells or Ly6a^Pos^ L-MCSs). They found that these cells express *Lumican* and *Ly6a* at a high level and therefore match clusters 2 and 6 in our study). The authors report that HYX exposure increases the number of these cells and alters their expression profile with the induction of pro-inflammatory, pro-fibrotic, and anti-angiogenic gene [39]. While this analysis describes the behavior of the rMC Sca1^Pos^ cells during acute injury, our study focusses on the status of these cells during the following “recovery” phase, as the animals are re-exposed to room air. Our analysis also allowed us to further refine the rMC Sca1^Pos^ population into 18 clusters, from which only 5 belong to the LIF cluster.

Due to the low number of rMC Sca1^Pos^ cells analyzed in the above mentioned study [39], it is challenging to state whether all the 18 clusters contained in the rMC Sca1^Pos^ cells are increased or if this increase is differentially impacting the different rMC Sca1^Pos^ clusters, with an increase in the MYF and ECM-producing cell clusters and a decrease in the LIF cluster. This is an essential aspect as only roughly 30% of the rMC Sca1^Pos^ cells, which correspond to the LIF cluster, display active niche activity [8].

In the future, a more detailed and powered analysis combining scRNAseq and functional assays of rMC Sca1^Pos^ cells at different time points during injury, recovery, and during FGF10 treatment will be crucial to better understand the impact of HYX on relevant rMC Sca1^Pos^ subpopulations.

## 5 Conclusion

In this study, we demonstrate that rFGF10 administration is able to induce de-novo alveologenesis in a BPD mouse model and identified subpopulations of rMC-Sca1^Pos^ niche cells potentially representing its cellular target.

## Acknowledgments (Funding)

This work was partially funded through Chiesi. S.B. was supported by grants from the Deutsche Forschungsgemeinschaft (DFG; BE4443/1-1, BE4443/4-1, BE4443/6-1, KFO309 P7 and SFB1213-projects A02 and A04) and DZL. S.H. was supported by the UKGM (FOKOOPV), the DZL, and University Hospital Giessen and grants from the DFG (KFO309 P2/8; SFB1021 C05, SFB TR84 B9).

## Author contributions

ST designed the study, carried out the experiments, analyzed the data, and wrote the manuscript. CMC recruited funding from Chiesi Pharmaceutical Company, contributed to study design, performed experiments and data analysis. JLG, LG, GM, SK, KG, JK, MH, AIVA contributed to performing experiments and data analysis. SH, CS, NW, FR, GA, LB, HE, PM provided feedback and helped shape the research and discussed the results, and contributed to the final manuscript. SB and SR supervised the progress of the project and the findings of this work, contributed to the interpretation of the results, and wrote the manuscript. All authors reviewed the results and contributed to the final manuscript.

## Data availability statement (Code availability for the analysis of scRNA-seq data)

All scRNA sequencing data, including raw fastq sequencing files, gene expression matrices, and associated cell metadata generated in this study, have been deposited in the NCBI’s Gene Expression Omnibus (GEO) database under: https://www.ncbi.nlm.nih.gov/geo/query/acc.cgi?acc=GSE190934

## Graphical abstract

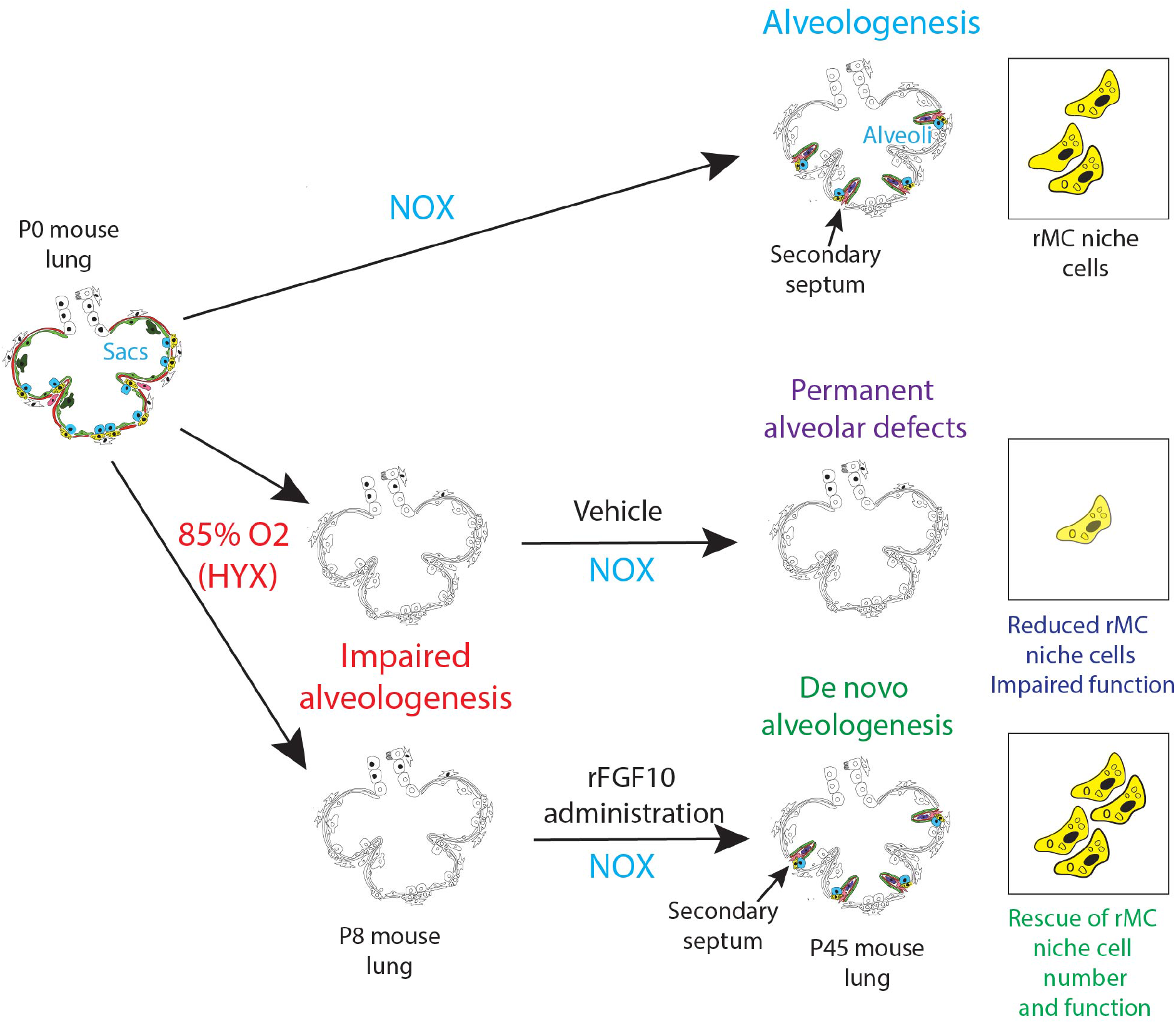

